# Age-Induced Changes in Mu Opioid Receptor Signaling in the Midbrain Periaqueductal Gray of Male and Female Rats

**DOI:** 10.1101/2022.01.13.475945

**Authors:** Evan F. Fullerton, Mary C. Karom, John M. Streicher, Larry J. Young, Anne Z. Murphy

## Abstract

**The analgesic effects of opioids are attenuated in aged rats.** Opioids such as morphine have decreased analgesic potency (but not efficacy) in aged rodents compared to adults; however, the neural mechanisms underlying this attenuated response are not yet known. The present study investigated the impact of advanced age and biological sex on opioid signaling in the ventrolateral periaqueductal gray (vlPAG) in the presence of chronic inflammatory pain. Assays measuring mu-opioid receptor (MOR) radioligand binding, GTPγS binding, receptor phosphorylation, cAMP inhibition, and regulator of G-protein signaling (RGS) protein expression were performed on vlPAG tissue from adult (2-3mos) and aged (16-18mos) male and female rats. Persistent inflammatory pain was induced by intraplantar injection of Complete Freund’s Adjuvant (CFA). Adult males exhibited the highest MOR binding potential and the highest G-protein activation (activation efficiency ratio) in comparison to aged males and females (adult and aged). No impact of advanced age or sex on MOR phosphorylation state was observed. DAMGO-induced cAMP inhibition was highest in the vlPAG of adult males compared to aged males and females (adult and aged). vlPAG levels of RGS4 and RGS9-2, critical for terminating G-protein signaling, were assessed using RNAscope. Adult rats (both males and females) exhibited lower levels of vlPAG RGS4 and RGS9-2 mRNA expression compared to aged males and females. The observed age-related reductions in vlPAG MOR binding potential, G-protein activation efficiency, and cAMP inhibition, along with the observed age-related increases in RGS4 and RGS9-2 vlPAG expression, provide potential mechanisms whereby the potency of opioids is decreased in the aged population. These results have significant implications for pain management in this population.

**Highlights:**

- Aged males and females (adult and aged) exhibit reduced vlPAG μ-opioid receptor binding potential compared to adult males.
- Aged males and females (adult and aged) exhibit reduced opioid-induced vlPAG G-protein activation compared to adult males.
- Aged males and females (adult and aged) exhibit reduced vlPAG MOR mediated cAMP inhibition compared to adult males.
- Aged rats (males and females) exhibit increased vlPAG mRNA expression of Regulator of G-Protein Signaling (RGS) proteins RGS4 and RGS9-2 compared to adult rats (males and females), which may explain the reduced receptor signaling observed in aged animals.
- These coordinate decreases in opioid receptor signaling may explain the previously reported reduced potency of opioids to produce pain relief in females and aged rats.

## 1 Introduction

Chronic pain is a debilitating condition that contributes to mental health disorders, disability, and increased risk of premature death (Domenichiello and Ramsden, 2019; Patel et al., 2013). More than 50% of individuals aged 65 and over suffer from chronic pain, and as the United States population rapidly ages, the development of effective strategies for chronic pain management for this demographic is imperative (Dahlhamer, 2018). Opioids, including morphine and fentanyl, are commonly prescribed analgesics for the management of chronic pain (Manchikanti et al., 2010; Sullivan et al., 2010), but effective dosing strategies for the elderly remain ill-defined, due in large part to a dearth of knowledge regarding the impact of advanced age on opioid signaling and analgesia.

Clinical studies examining pain management in the elderly are challenging; co-morbid conditions such as diabetes and high blood pressure, and participant use of concomitant medications affect patient outcomes, contributing to difficulties in the interpretation of results (Naples et al., 2016; Prostran et al., 2016). Additionally, there exist age- and sex-related individual differences in the likelihood of reporting pain in a clinical setting which may lead to misrepresentations of analgesic efficacy (Dampier et al., 2013; Reddy et al., 2012). Particularly, aged populations are known to underreport pain due to fears of institutionalization and concerns with addiction and overdose (Reddy et al., 2012).

Using a preclinical model of persistent inflammatory pain (intraplantar Complete Freund’s Adjuvant; (CFA)), we have previously reported a significant impact of age and sex on morphine potency. Specifically, we showed that aged rats (18mos) exhibit a decreased morphine potency compared to adults (2mos), with aged males requiring greater than 2x the concentration of morphine than their adult counterparts to produce equivalent analgesia (Fullerton et al., 2021). Similar results have been reported in prior preclinical studies, suggesting that aged rodents require higher doses of opioids to produce antinociception (Kavaliers et al., 1983; Kramer and Bodnar, 1986; Webster et al., 1976). The mechanisms underlying the attenuation of opioid potency in the aged population are currently unknown.

Morphine-induced analgesia is mediated primarily via binding to μ-opioid receptors (MOR), prototypical G-protein coupled receptors (GPCRs) located predominantly on neuronal cell membranes (Martin, 1963; Wolozin and Pasternak, 1981). Following agonist binding, coupled G-proteins undergo a conformational change in which the guanosine diphosphate (GDP) bound to the inactive alpha subunit is replaced by guanosine triphosphate (GTP), activating the subunit and promoting opioid signaling through interaction with downstream effectors (Senese et al., 2020). MORs are coupled to a family of G-proteins called Gi/o that signal via inhibition of adenylyl cyclase and subsequently decrease cyclic adenosine monophosphate (cAMP) (Koehl et al., 2018; Bouchet et al., 2021). This G-protein signaling is downregulated or terminated by **R**egulator of **G**-Protein **S**ignaling (RGS) proteins, which act as GTPase activating proteins (GAPS), hydrolyzing the active GTP back into GDP and terminating downstream signaling (Gerber et al., 2016). RGS proteins, particularly RGS4 and RGS9-2, have been previously implicated in opioid-mediated G-protein signaling, and have been shown to attenuate analgesia (Garnier et al., 2003; Psifogeorgou et al., 2007; Avrampou et al., 2019; Senese et al., 2020). A primary target of MOR downstream signaling is the effector adenylate cyclase (AC). AC enzymatically converts adenosine triphosphate (ATP) to the excitatory second messenger cyclic adenosine monophosphate (cAMP). MOR binding inhibits AC, thereby reducing cAMP expression, facilitating neuronal hyperpolarization, and promoting analgesia (Santhappan et al., 2015). Like other G-protein coupled receptors, MOR signaling is subject to desensitization via the cellular mechanism of receptor phosphorylation, which limits agonist binding and recruits arrestin proteins, and thus attenuates opioid signaling. (Zhang et al., 1996; Yu et al., 1997; Groer et al., 2011).

The present studies test the hypothesis that the observed age-induced reduction in morphine potency in the presence of chronic pain is mediated by changes in MOR signaling within the midbrain periaqueductal gray, a CNS region critical for the opioid modulation of pain. (Basbaum et al., 1976; Behbehani and Fields, 1979; Morgan et al., 2006, 1992). The ventrolateral PAG (vlPAG) contains a large population of MOR+ neurons, and direct administration of MOR agonists into the PAG produces potent analgesia (Satoh et al., 1983; Jensen and Yaksh, 1986; Bodnar et al., 1988; Loyd et al., 2008) In addition, intra-PAG administration of MOR antagonists or lesions of PAG MOR significantly attenuate the analgesic effect of systemic morphine, suggesting a critical role for PAG MOR in mediating morphine action (Ma and Han, 1991; Y. Zhang et al., 1998; Loyd et al., 2008).

We have previously shown that aged rats exhibit reduced vlPAG MOR protein expression and reduced MOR agonist binding in the vlPAG compared to adult rats (Fullerton et al., 2021), suggesting that diminished levels of functioning MOR in the vlPAG contribute to the attenuated opioid potency seen in the aged. The present studies build on these previous findings to assess age and sex differences in MOR availability, ligand affinity, phosphorylation, G-protein activation, cAMP inhibition, and expression of RGS proteins. We report that aged rats, and more specifically males, exhibit reduced MOR binding potential, reduced G-protein activation, reduced cAMP inhibition, and increased RGS expression compared to adults, with no change in MOR phosphorylation. Similar changes were observed in females regardless of age. These findings identify several mechanisms whereby opioid signaling is attenuated in the aged rat, and provide a framework for the development of novel pharmacological therapies to improve pain management in the elderly.

## 2 Materials and methods

### 2.1 Experimental subjects

Adult (2-3mos) and aged (16-18mos) male and regularly cycling female Sprague–Dawley rats were used in these experiments (Charles River Laboratories, Boston, MA). Rats were co-housed in same-sex pairs on a 12:12 hour light/dark cycle (lights on at 08:00 am). Access to food and water was ad libitum throughout the experiment, except during testing. All studies were approved by the Institutional Animal Care and Use Committee at Georgia State University and performed in compliance with Ethical Issues of the International Association for the Study of Pain and National Institutes of Health. All efforts were made to reduce the number of rats used in these experiments and to minimize pain and suffering. All assays were performed on separate cohorts of rats (n=8/sex/age; N=32), with the exception of the radioligand binding and GTPγS assays, which were run simultaneously on PAG tissue from a single cohort.

### 2.2 Vaginal cytology

Beginning ten days prior to testing, vaginal lavages were performed daily on adult and aged female rats to confirm that all rats were cycling regularly and to keep daily records of the stage of estrous. Proestrus was identified as a predominance of nucleated epithelial cells, and estrus was identified as a predominance of cornified epithelial cells. Diestrus 1 was differentiated from diestrus 2 by the presence of leukocytes. Rats that appeared between phases were noted as being in the more advanced stage (Loyd et al., 2007).

### 2.3 CFA-induced chronic pain treatment

72 hours prior to experimentation, persistent inflammatory pain was induced by injection of complete Freund’s adjuvant (CFA; Mycobacterium tuberculosis; Sigma; 200μl), suspended in an oil/saline (1:1) emulsion, into the plantar surface of the right hind paw. Edema was present within 24 hours of injection, indicated by a >100% change in paw diameter, determined using calibrated calipers applied midpoint across the plantar surface compared to handled paw.

### 2.4 Membrane preparation for radioligand binding and GTPγS assays

vlPAG membrane protein lysates were prepared from adult and aged, male and female, handled and CFA treated rats to be used for radioligand binding and GTPγS assays. 72 hours post-CFA injection or handling, rats were restrained using DecapiCones and decapitated. Brains were removed rapidly, flash-frozen in 2-methyl butane on dry ice, and stored at −80°C. Ventrolateral PAG tissue from caudal PAG (Bregma −7.5 through −8.5) was dissected from each brain using a straight edge razor at −20°C. On the day of the assay, PAG sections were placed in ice-cold assay buffer (50 mM Tris-HCl, pH 7.4). Tissue was homogenized with a glass dounce and centrifuged at 20,000g at 4°C for 30 min. The supernatant was discarded and the pellet resuspended in assay buffer (Li et al., 2018; Shaqura et al., 2016b, 2016a; Zollner et al., 2003). Membrane protein concentration was calculated using the Bradford Assay, and lysates of vlPAG membrane protein from adult and aged, male and female, naïve and CFA treated rats (n=4; N=32) were used immediately for radioligand binding and GTPγS assays.

### 2.5 Saturation radioligand binding assay

Saturation binding experiments were performed on vlPAG membranes using [^3^H]DAMGO (specific activity 50 Ci/mmol, American Radiolabeled Chemicals, Missouri). Briefly, 100 µg of membrane protein was incubated with various concentrations of [^3^H]DAMGO (0.5 to 5nM), in a total volume of 1ml of binding buffer (50mM Tris-HCl pH 7.4). Nonspecific binding was defined as radioactivity remaining bound in the presence of 10 µM unlabeled naloxone. At the end of the incubation period (1h at RT) bound and free ligands were separated by rapid filtration over Whatman brand Grade GF/C glass filters (Sigma-Aldrich) using a sampling vacuum manifold (MilliporeSigma). Filters were washed four times with 5ml of cold dH_2_0. Bound radioactivity was determined by liquid scintillation spectrophotometry after overnight extraction of the filters in 3ml of scintillation fluid. Specific activity was calculated by subtracting the non-specific binding mean value at each concentration of [^3^H]DAMGO. All experiments were performed in triplicate. B_max_ and K_d_ values were determined by nonlinear regression analysis of concentration-effect curve in GraphPad Prism 9.1.

### 2.6 Agonist-stimulated [^35^S]GTPγS binding

DAMGO-stimulated [^35^S]GTPγS binding to PAG membrane protein was assessed by incubating membrane protein (100ug) in the presence or absence of [^35^S]GTPγS (0.1nM) (specific activity 1250 Ci/mmol, American Radiolabeled Chemicals, Missouri) and various concentrations of DAMGO (2 to 30,000 nM) in assay buffer (20mM Tris, 10mM MgCl_2_, 100mM NaCl, 0.2mM EGTA, pH 7.4) for 30min at 30°C. Stimulated [^35^S]GTPγS binding was compared to unstimulated binding at each measurement point and presented as percent basal binding. At the end of the incubation period, bound and free ligands were separated by rapid filtration over Whatman brand Grade GF/B glass filters (Sigma-Aldrich) using a sampling vacuum manifold (MilliporeSigma). Filters were washed four times with 5ml of cold buffer (50 mM Tris-HCl, pH 7.4). Bound radioactivity was determined by liquid scintillation spectrophotometry after overnight extraction of the filters in 3ml of scintillation fluid. All experiments were performed in duplicate. Efficacy (Emax) is defined as the maximum percent stimulation by an agonist; potency (EC_50_) is defined as the concentration of DAMGO required for half the maximal response. Emax and EC_50_ values were determined by nonlinear regression analysis of concentration-effect curves using Graph Pad Prism 9.1.

### 2.7 Phosphorylated MOR analysis

To assess the impact of advanced age and sex on MOR phosphorylation state, levels of MOR phosphorylation were analyzed by Western blot. The lysis buffers used and general methods are the same as reported in (Lei et al., 2017). Briefly, rat PAG samples were homogenized in ∼500 μL of lysis buffer using a glass dounce and rotated overnight at 4°C. The samples were then spun down at 13,000g for 10 min at 4°C. The resulting lysates were quantified by DC Protein Assay Kit (Bio-Rad, #500-0111) according to the manufacturer’s instructions. 50-75 μg of soluble protein per sample (same for all samples in a set) was run on 10% Bis-Tris Bolt PAGE gels (Fisher Scientific, #NW00100BOX), and wet-transferred to nitrocellulose membrane (Protran 0.2 μm NC, #45-004-001 from Fisher Scientific) at 30V at 4°C for 2-3 hours. Membranes were blocked using 5% nonfat dry milk in TBS, then blotted for primary antibody target in 5% BSA in TBST overnight rocking at 4°C. Primary antibodies used were: pMOR (1:1000, Bioss #bs-3724R), tMOR (1 μg/mL, R&D #MAB6866), and GAPDH (1:1000, Invitrogen #MA5-15738). All 3 targets were always probed for on the same blot, using low pH stripping buffer between each set. Secondary antibodies were used at 1:5000 each in 5% nonfat dry milk in TBST, rocking for ∼1 hr at room temperature. Secondary antibodies used were Goat-α-Mouse-IRDye800CW (Fisher Scientific #NC9401841) and Goat-α-Mouse-IRDye680LT (Fisher Scientific #NC0046410). Target signal was acquired using an Azure Sapphire imager using the near-infrared channels (658 and 784 nm). Band density was quantified, and background subtracted using the onboard AzureSpot analysis software. Samples were run with at least two representatives of each experimental group on the same gel. Adult and aged, male and female, naïve and CFA treated rats (n=4; N=32) were used for these experiments. pMOR signal was normalized to the same sample tMOR or GAPDH as indicated, and all samples were normalized to the Adult Male average on each gel before combining data from different gels.

### 2.8 Agonist-dependent cAMP inhibition

DAMGO-dependent cAMP inhibition was analyzed using a cAMP assay. vlPAG membrane protein lysates were prepared from adult and aged, male and female, handled and CFA treated rats. 72 hours post-CFA injection or handling, rats were restrained using DecapiCones and decapitated. Brains were removed rapidly, flash-frozen in 2-methyl butane on dry ice, and stored at −80°C. Ventrolateral PAG tissue from caudal PAG (Bregma −7.5 through −8.5) was dissected from each brain using a straight edge razor at −20°C. Briefly, vlPAG tissues were homogenized in ice-cold lysis buffer containing 0.25-M sucrose, 50-mM Tris–HCl pH 7.5, 5-mM EGTA, 5-mM EDTA, 1-mM phenylmethylsulfonyl fluoride, 0.1-mM dithiothreitol, and 10 μg/ml leupeptin. The homogenized tissues were then centrifuged at 1000×*g* for 5 min (4 °C) and the supernatant was centrifuged at 35000 ×*g* for 10 min (Viganò et al., 2003). vlPAG membrane protein samples (100ug) from adult and aged, male and female, naïve and CFA treated rats (n=4; N=32) were incubated in 1uM forskolin (FSK) and in the presence or absence of 10uM DAMGO to stimulate adenylate cyclase activity. cAMP levels in vlPAG tissue were determined using the LANCE *Ultra* cAMP Detection Kit (Perkin Elmer) a time-resolved fluorescence resonance energy transfer (TR-FRET) cAMP immunoassay. Data are plotted as % change in TR-FRET signal comparing FSK baseline measurements to measurements following DAMGO-stimulated cAMP inhibition.

### 2.9 Single-molecule fluorescence in situ hybridization

Single-molecule fluorescent in situ hybridization (RNAscope) assays were used to determine mRNA expression of OPRM1, RGS9-2, and RGS4 in the vlPAG. Rats were restrained using DecapiCones and decapitated. Brains were removed rapidly, flash frozen in 2-methyl butane on dry ice, and stored RNase free at −80 °C. Frozen tissue was sectioned in a 1:6 series of 20 μm coronal sections at –20 °C with a Leica CM3050S cryostat. Sections were immediately mounted onto Superfrost slides (20 °C) and stored at −80 °C until the time of the assay. vlPAG sections from adult and aged, male and female, naïve and morphine treated rats were used (n=4; N=32). For morphine dosing paradigm see (Fullerton et al., 2021). Tissue was processed for single-molecule fluorescence in situ hybridization (smFISH) according to the RNAscope Multiplex Kit protocol (Advanced Cell Diagnostics) using probes for OPRM1, RGS4, and RGS9-2. To facilitate cellular mRNA quantification within the vlPAG, sections were counterstained with DAPI. mRNA puncta were visualized as fluorescent signal. Fluorescent images were captured on Zeiss LSM 700 Confocal Microscope at 40x, and mRNA expression (target puncta/DAPI) was calculated using Imaris software. To determine RGS4 and RGS9-2 expression levels in MOR+ neurons, quantification was restricted to puncta located within 10μm of DAPI that co-expressed OPRM1 mRNA. mRNA expression values were determined for the left and right ventrolateral subdivisions of each PAG image from two representative levels of the mid-caudal PAG (Bregma −7.74 and −8.00). As there was no significant effect of rostrocaudal level in the analyses, data were collapsed and presented as vlPAG and were averaged for each rat. RGS4 and RGS9-2 mRNA values are expressed as the mean ± standard error of the mean (SEM). All images were collected and analyzed by an experimenter blinded to the experimental condition.

### 2.10 Statistical analysis and data presentation

All values are reported as mean ± SEM. Data were assessed for normality and homogeneity of variance using Shapiro-Wilk and Bartlett’s tests. Significant main effects of sex, age, and treatment were assessed using ANOVA; p < 0.05 was considered statistically significant. Tukey’s post-hoc tests were conducted to determine significant mean differences between groups that were a priori specified. Data are expressed either as fmol/protein or nM DAMGO for radioligand binding assay, or Emax or EC50 for GTPγS assay.

## 3 Results

### 3.1 Advanced age and sex impact vlPAG MOR binding properties

To assess the impact of advanced age, biological sex, and pain on vlPAG MOR signaling, we first used radioligand saturation binding assays to determine MOR binding parameters. The saturation binding curves generated with [^3^H]DAMGO are shown in Fig. 1A. There was no significant impact of CFA treatment on any measures, so CFA and handled groups were combined. Kd values, indicative of receptor affinity, were determined using the concentration-effect curves generated from each sample. Analysis of Kd values indicated no significant impact of age [F_(1,26)_ = 0.770, p=0.388] or sex [F_(1,26)_ = 1.241, p=0.275], or a significant interaction [F_(1,26)_ = 1.278, p=0.269] (Fig. 1B).

**Fig. 1.**
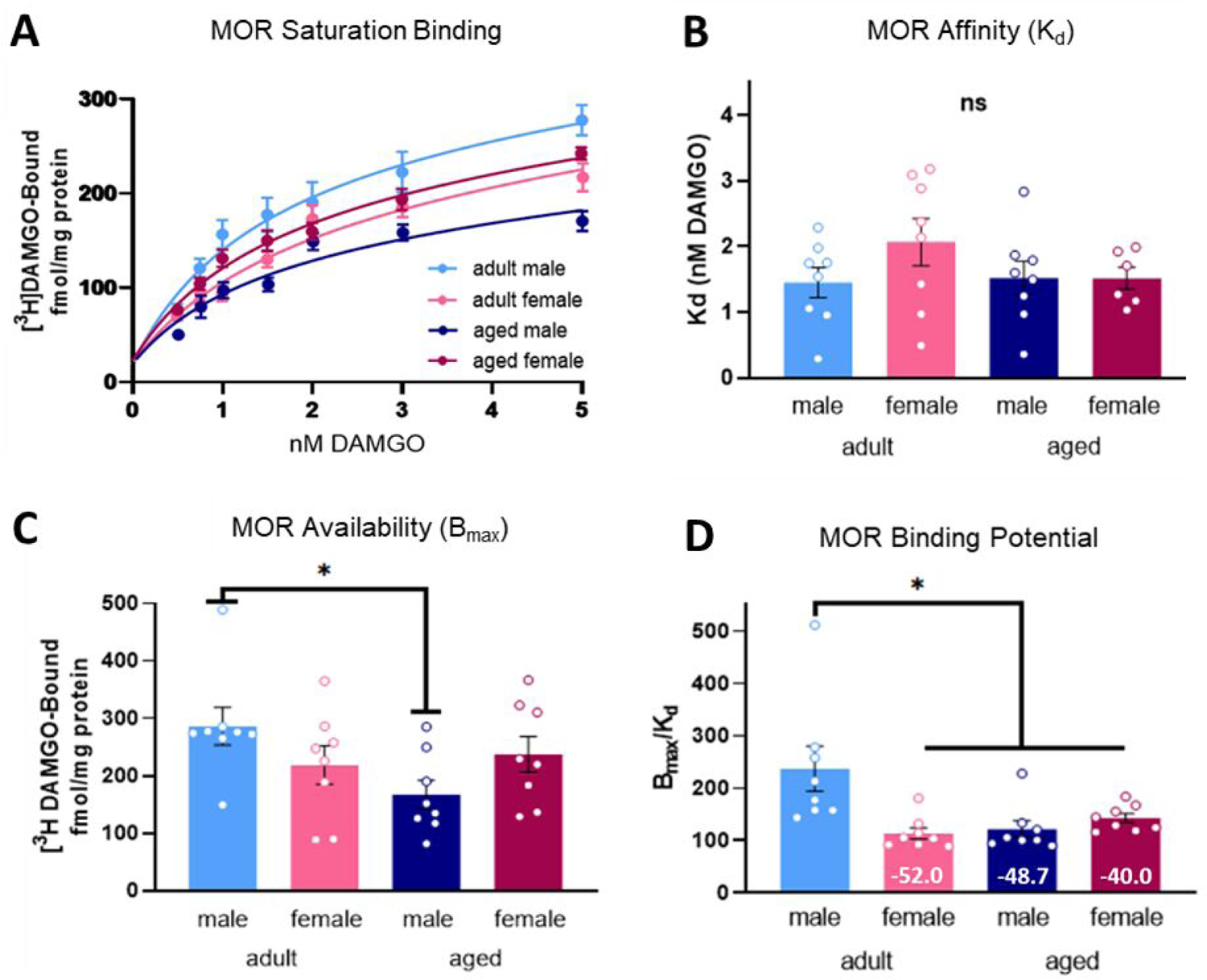
Saturation binding curve of bound [3H]DAMGO. (A). No significant effect of age or sex was observed in MOR affinity (K_d_) (B). MOR availability (B_max_) was significantly lower in aged males compared to adult males (C). Adult females, aged males, and aged females exhibited reduced MOR binding potential (B_max_/K_d_) compared to adult males, indicating attenuated agonist interaction at the receptor level (D). ns, not significant. *Significant difference between adult and aged males, or adult males and adult females; p<0.05 calculated by Tukey’s post hoc test. Graphs indicate mean ± SEM. Values indicate % change from adult male.

Analyses of B_max_ values, indicative of receptor availability, showed no significant impact of age [F_(1,28)_ = 2.661, p=0.114] or sex [F_(1,28)_ = 0.002, p=0.967], but a significant interaction [F_(1,28)_ = 4.995, p=0.034]. Post hoc analysis showed a significant difference in vlPAG MOR availability between adult males and aged males, as evidenced by reduced B_max_ values in the aged males compared to their adult counterparts (p = 0.025)(Fig. 1C). B_max_ values for aged males were markedly reduced compared to adult males, while adult females and aged females exhibited smaller reductions in B_max_ compared to their adult male counterparts (%Δ −23.6 and −17.1, respectively. Fig. 1C).

Analyses of binding potential (BP) values, a measure that takes into consideration the density of available receptors and the affinity of the receptor for its agonist, showed a significant impact of sex, with males exhibiting greater binding potential than females [F_(1,28)_ = 4.631, p=0.040] and a significant interaction between age and sex [F_(1,28)_ = 9.110, p=0.005]. No significant impact of age [F_(1,28)_ = 3.312, p=0.079] was observed. Post hoc analyses showed a significant difference in vlPAG MOR binding potential between adult males and adult females (p = 0.006), and adult males and aged males (p = 0.010), indicating that adult males exhibit greater vlPAG MOR binding potential than their aged male counterparts and their adult female counterparts (Fig. 1D). Adult females, aged males, and aged females all exhibited marked reductions in MOR binding potential compared to their adult male counterparts (%Δ −52, −48.7, and −40, respectively; Fig. 1D).

### 3.2 Advanced age and sex impact opioid-induced G-protein activation in the vlPAG

We next conducted GTPγS binding assays to determine if advanced age or biological sex impacted MOR mediated G-protein activation. Preliminary studies revealed a significant main effect of chronic pain [F_(1,23)_ = 21.16, p = 0.0001]. To improve the translatability of these results to the target population of aged patients suffering from chronic pain, GTPγS experiments were performed exclusively on CFA-treated rats. Concentration curves generated from [^35^S]GTPγS assays are shown in Fig. 2A. Analyses of E_max_ values, a measure of G-protein availability combined with ligand efficacy, indicated no significant impact of age [F_(1,24)_ = 0.035, p = 0.8532] or sex [F_(1,25)_ = 2.852, p = 0.1042], or a significant interaction [F_(1,24)_ = 2.023, p = 0.1678] (Fig. 2B).

**Fig 2.**
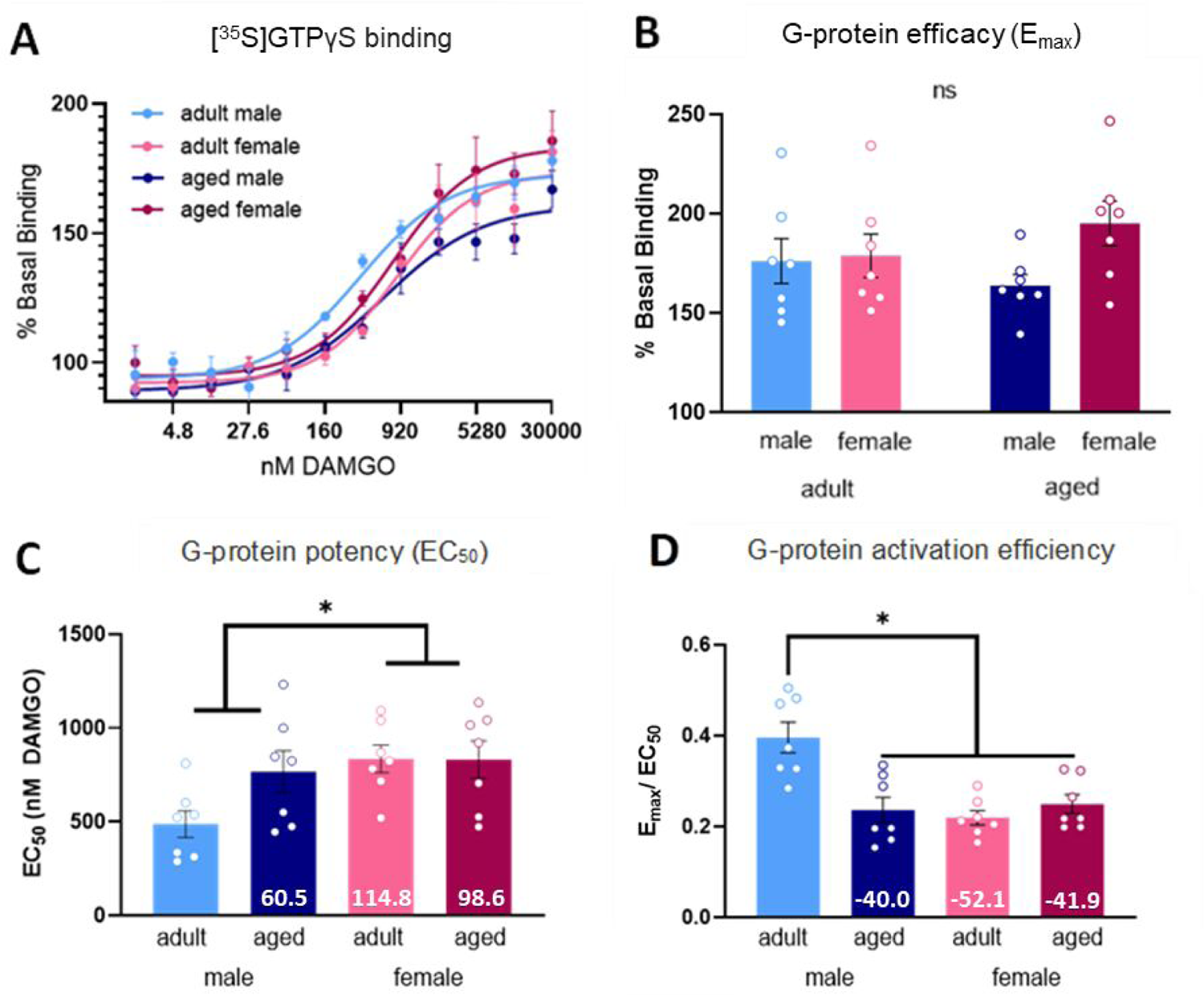
[^35^S]GTPγS binding curve of agonist-stimulated GTPγS binding. (A). No significant effect of age or sex was observed in E_max_ (B). Aged males, adult females, and aged females exhibited increased EC_50_ values compared to adult males, indicating that adult males have the greatest potency (C). Aged males, adult females, and aged females exhibited reduced G-protein activation efficiency (E_max_/EC_50_) compared to adult males, indicating attenuated G-protein signaling (D). ns, not significant. *Significant difference between adult and aged males, or adult males and adult females; p<0.05 calculated by Tukey’s post hoc test. Graphs indicate mean ± SEM. Values indicate % change from adult male.

Analyses of EC_50_ values, a measure of the concentration of DAMGO required for half-maximal G-protein binding, indicated a significant impact of sex, with males exhibiting lower effective concentration values than females, reflecting a higher potency of activation [F_(1,24)_ = 5.177, p = 0.0321]. There was no significant impact of age [F_(1,24)_ = 2.339, p = 0.1393], and no significant interaction [F_(1,24)_ = 2.473, p = 0.1289] (Fig. 2C). Compared to their adult male counterparts, aged males, adult females, and aged females all exhibited marked increases in G-protein EC_50_ as evidenced by their percent change values (%Δ 60.5, 114.8, and 98.6, respectively), reflecting a lower potency of activation (Fig. 2C). Together, these results suggest that opioid-induced G-protein signaling is impaired in the PAG of the aged rat and the female rat.

Next, we calculated the coefficient ratio of analysis of E_max_ / EC_50_, which is indicative of overall G-protein activation efficiency. A significant impact of age was noted, with adults exhibiting higher vlPAG G-protein activation efficiency than aged rats [F_(1,24)_ = 6.491, p = 0.0177]. There was also a significant impact of sex, with males exhibiting higher vlPAG G-protein activation efficiency compared to females [F_(1,24)_ =10.39, p = 0.0036]. A significant interaction was also observed [F_(1,24)_ = 13.93, p = 0.0010]. Post hoc analyses showed a significant difference in vlPAG G-protein activation efficiency between adult males and aged males (p = 0.0009), with adult males exhibiting greater vlPAG G-protein activation efficiency than aged males. A significant difference in vlPAG G-protein activation efficiency between adult males and adult females (p= 0.0003) was also observed, with adult males exhibiting greater vlPAG G-protein activation efficiency than adult females (Fig. 2D). Aged males, adult females, and aged females all exhibited marked reductions in G-protein activation efficiency compared to their adult male counterparts (%Δ −40.0, −52.1, and −41.9, respectively; Fig. 2D).

### 3.3 Advanced age and sex do not impact phosphorylated MOR in the vlPAG

The results above showed significant reductions in MOR binding potential and G-protein activation efficacy. Therefore, we next examined if there was an effect of age and sex on MOR phosphorylation, as increased levels of pMOR would likely contribute to these reductions. Western blots were used to determine the impact of advanced age on MOR phosphorylation at serine-375, a known site of phosphorylation-mediated desensitization (Schulz et al., 2004). There was no significant impact of treatment, so CFA and handled groups were combined. Although increased pMOR/tMOR was observed in aged males compared to their adult counterparts (%Δ 34.0), no significant main effect of age [F_(1,41)_ = 2.168, p = 0.1486], sex [F_(1,41)_ = 1.807, p = 0.1863], or age x sex interaction [F_(1,41)_ = 0.4615, p = 0.5008] was observed (Fig. 3).

**Fig 3.**
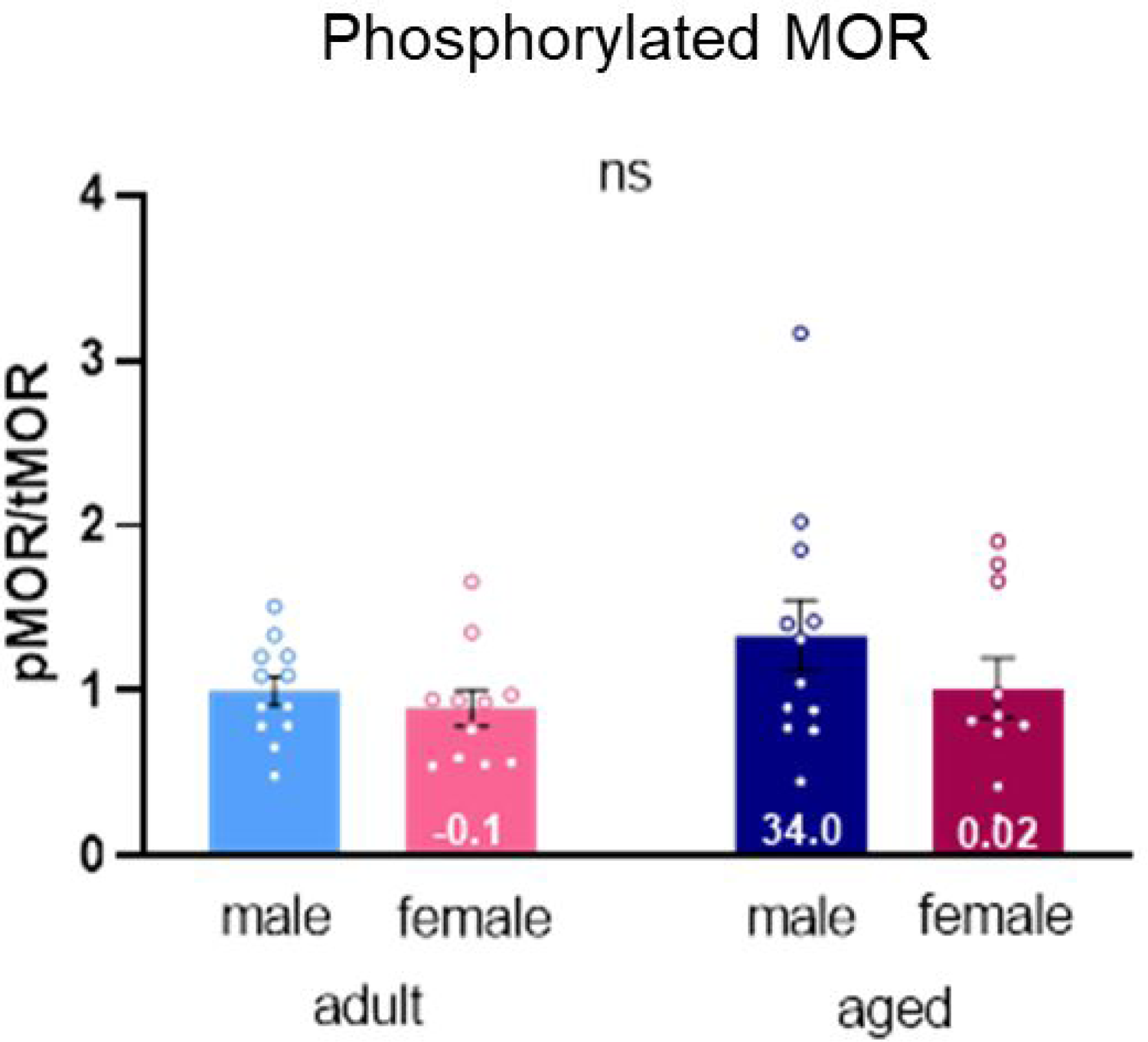
No significant impact of advanced age or biological sex on the ratio of phosphorylated MOR/ total MOR. ns, not significant. Graph indicates mean ± SEM. Values indicate % change from adult male.

### 3.4 Advanced age and sex impact opioid-induced cAMP inhibition in the vlPAG

We next determined if advanced age, biological sex, or persistent pain impacted opioid-induced cAMP inhibition in the vlPAG. Forskolin-induced cAMP release was used as a baseline measurement for each group, while forskolin + DAMGO was used to assess the degree to which cAMP was inhibited by DAMGO. Percent change from baseline was used to compare across treatment groups (Fig. 4A). There was no significant impact of persistent pain on any measures, so CFA and handled groups were combined. Our analyses revealed a significant main effect of age [F_(1,30)_ = 9.314, p = 0.0047], a significant main effect of sex [F_(1,30)_ = 10.31, p = 0.0031], and a significant interaction [F_(1,30)_ = 14.24, p = 0.0007].

**Fig 4.**
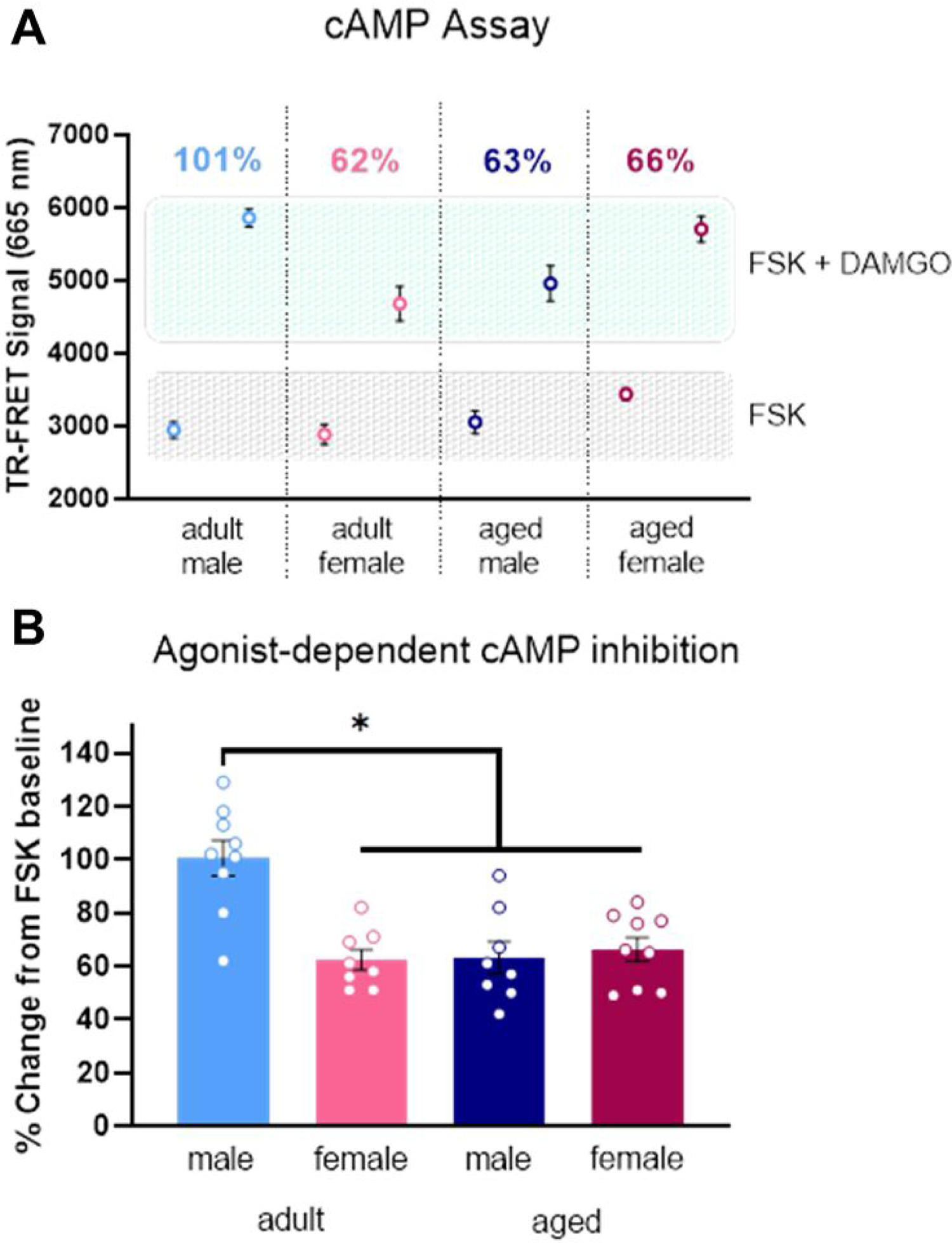
TR-FRET immunoassay showing % change from FSK stimulated baseline and DAMGO inhibited cAMP (cAMP levels are inversely proportional to TR-FRET signal). Adult males exhibited the highest level of agonist-dependent % change in cAMP (A). Aged males and females exhibited reduced agonist-dependent cAMP inhibition compared to adult males, indicating attenuated downstream opioid signaling (B). *Significant difference between adult and aged males, or adult males and adult females; p<0.05 calculated by Tukey’s post hoc test. Graphs indicate mean ± SEM. Values indicate % change from FSK baseline.

Post hoc analyses showed a significant difference in cAMP inhibition between adult males and adult females (p = 0.0002), adult males and aged males (p = 0.002), and adult males and aged females (p = 0.004), indicating that DAMGO elicits greater cAMP inhibition in adult males compared to aged males and females (both adult and aged) (Fig. 4B).

### 3.5 Advanced age and sex impact RGS4 and RGS9-2 expression in the vlPAG

Regulator of G-protein Signaling (RGS) proteins act as GAP accelerators to negatively modulate G-protein signaling. RGS protein family members RGS4 and RGS9-2 are expressed in the vlPAG and have both been shown to regulate opioid signaling by reversing G-protein activation. Therefore, we next used smFISH to determine if RGS4 and RGS9-2 expression in the vlPAG was altered by advanced age and biological sex. In these studies, following CFA administration, a cohort of rats was administered morphine to examine the relationship between morphine EC_50_ and RGS levels. However, no differences were noted between the morphine and naive conditions, so these groups were combined. First, we assessed total vlPAG expression of RGS4 and RGS9-2. These analyses revealed a significant main effect of age on both RGS4 [F_(1,37)_ = 18.15, p = 0.0001] and RGS9-2 [F_(1,41)_ = 17.09, p < 0.0002], with aged rats exhibiting increased levels of RGS4 and RGS9-2 mRNA compared to their adult counterparts (Fig. 5A & B). No significant main effect of sex on RGS4 [F_(1,37)_ = 0.0247, p = 0.8763] or RGS9-2 [F_(1,41)_ = 0.4855, p = 0.4903], or significant interactions between age and sex for RGS4 [F_(1,37)_ = 0.4994, p = 0.4842] or RGS9-2 [F_(1,41)_ = 0.3246, p = 0.5723] were observed (Fig. 5A & B).

**Fig 5.**
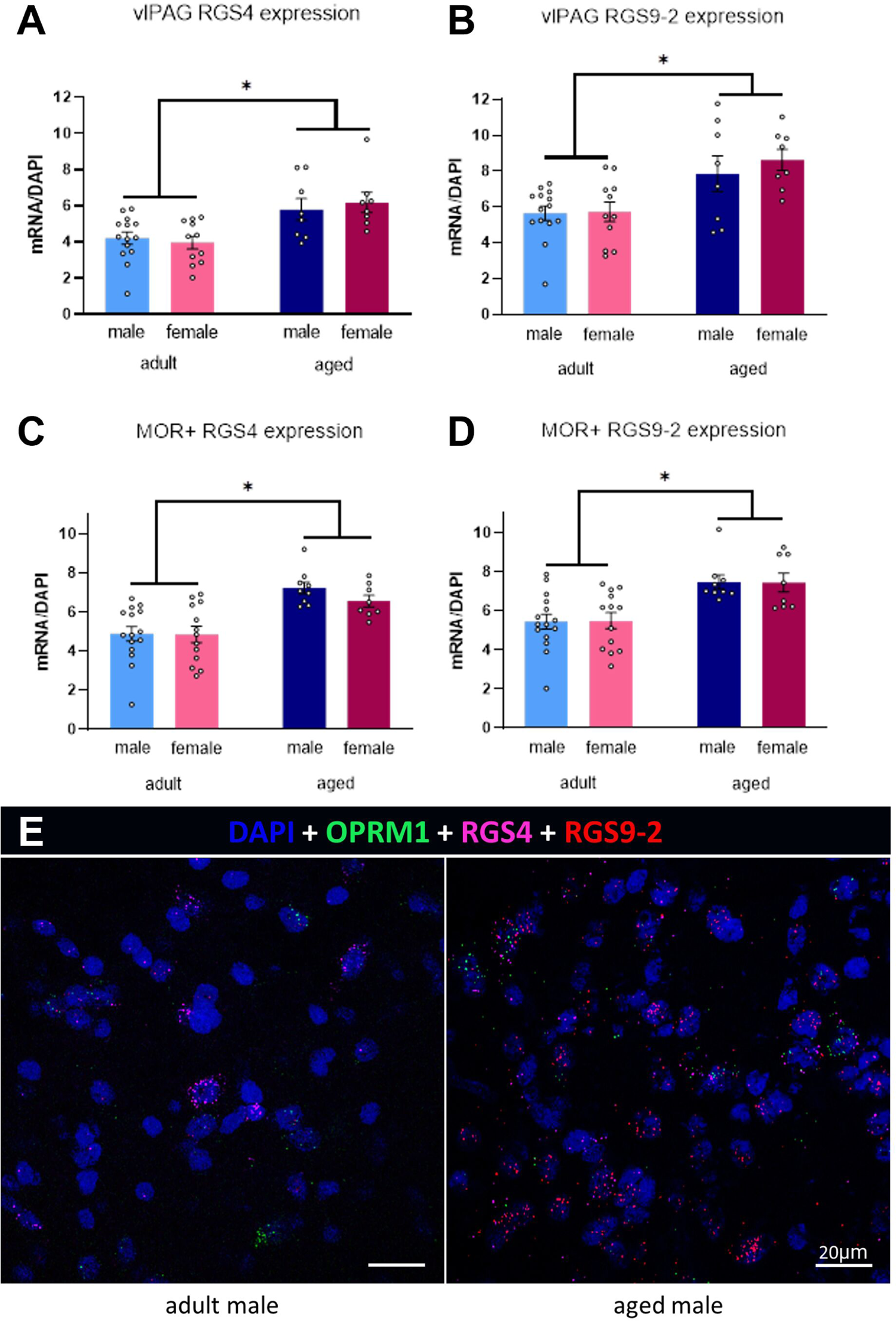
mRNA expression of RGS4 (A) and RGS9-2 (B) in the vlPAG was significantly greater in aged rats compared to adults. mRNA expression of RGS4 (C) and RGS9-2 (D) on MOR+ cells in the vlPAG was significantly greater in aged rats compared to adults. (E) Photomicrograph of RGS expression compared an adult male and an aged male PAG section. *Significant difference between adults and aged rats; p<0.05 calculated by 2×2 ANOVA. Graphs indicate mean ± SEM.

Following our assessment of total RGS4 and RGS9-2, we next restricted our analyses to RGS4 and RGS9-2 mRNA expressed on MOR+ neurons. Similar to what was noted above, a significant main effect of age on expression of RGS4 [F_(1,41)_ = 26.47, p < 0.0001] and RGS9-2 [F_(1,41)_ = 21.69, p < 0.0001] was observed, with aged rats exhibiting increased levels of RGS4 and RGS9-2 mRNA in MOR+ neurons compared to their adult counterparts (Fig. 5C & D). No main effect of sex for RGS4 [F_(1,41)_ = 0.7881, p = 0.3799] or RGS9-2 [F_(1,41)_ = 0.0006, p = 0.9799], or significant interactions between age and sex for RGS4 [F_(1,41)_ = 0.6583, p = 0.4219] or RGS9-2 [F_(1,41)_ = 0.0075, p = 0.9314] were observed (Fig. 5C & D).

## 4 Discussion

The present studies are the first to show that advanced age results in a significant attenuation in MOR signaling within the vlPAG of male and female rats. Specifically, aged males and females (regardless of age) showed decreased MOR binding potential, decreased G-protein activation, and decreased agonist-stimulated cAMP inhibition in comparison to adult males (Fig. 6). These changes, along with the observed increase in RGS4 and RGS9-2 expression, provide a mechanism whereby morphine potency is significantly reduced in aged rats (Fullerton et al., 2021). These changes also support our earlier observed sex difference whereby adult females have a lower opioid analgesic potency than adult males, equivalent to aged males and females.

**Fig 6.**
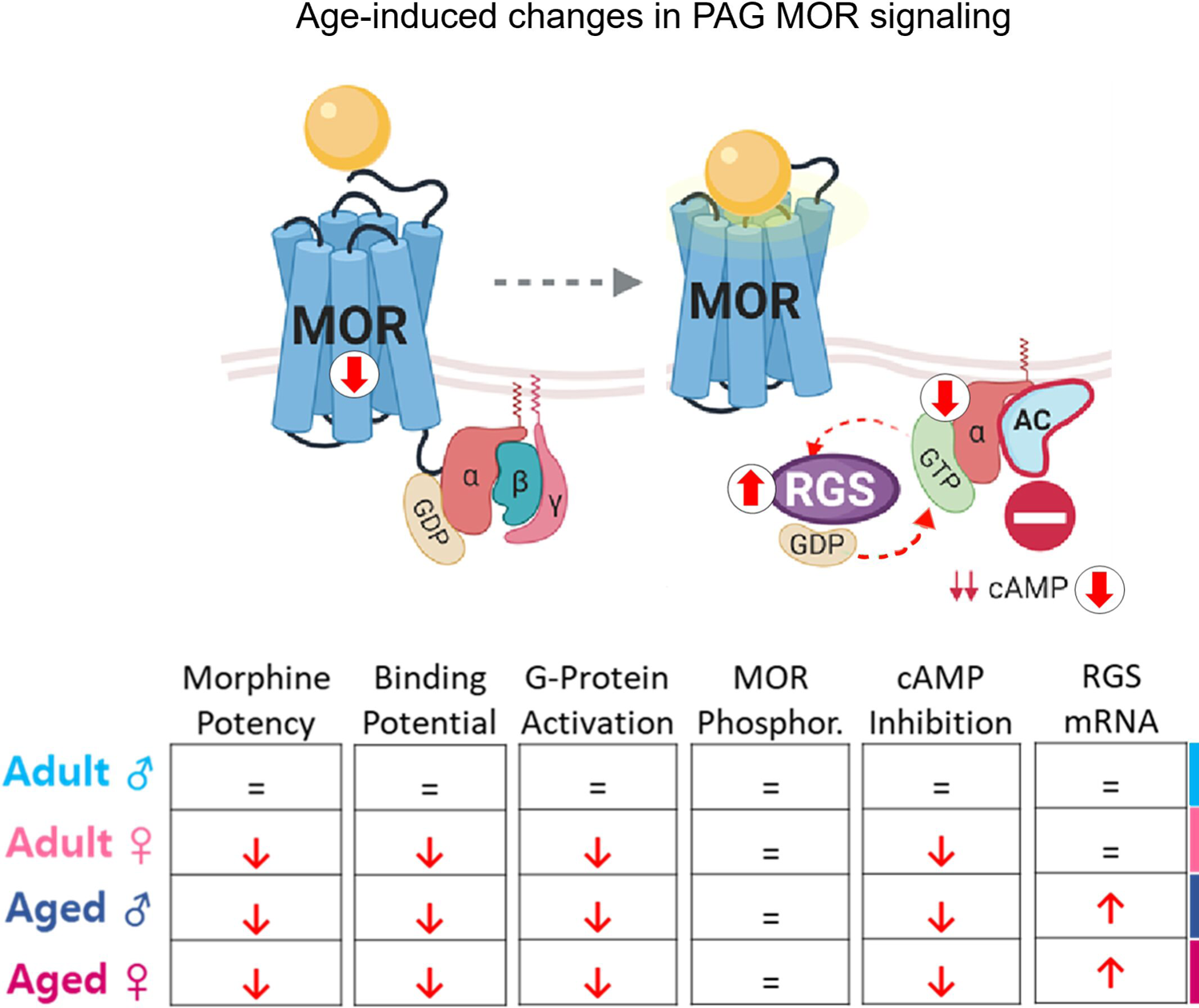
Model and table of MOR signaling impairments in the vlPAG of the aged and female rat. Aged and female rats exhibit decreased MOR binding potential, decreased G-protein activation, and decreased cAMP inhibition, while aged rats alone demonstrate increased RGS expression. No impact of age or sex was found in phosphorylation of MOR at serine-375

### Impact of advanced age and sex on MOR binding potential

In our previous study, we reported that morphine potency to block persistent inflammatory pain is reduced in aged and female rats, likely due to a reduction in DAMGO binding in the vlPAG (Fullerton et al., 2021). Consistent with that finding, we report here that the vlPAG MOR binding potential in aged males and females of either age is significantly reduced in comparison to adult male rats, providing additional support that the decrease in opioid potency observed in these groups is driven, in part, by reduced expression/availability of MOR in the vlPAG. Notably, we did not observe an impact of advanced age or sex on the MOR’s affinity for its ligand, as there were no significant effects of age or sex on DAMGO K_d_ values. However, aged males had reduced MOR availability as evidenced by lower B_max_ values compared to their adult male counterparts. This result is consistent with our previous finding that aged males exhibited reduced DAMGO binding in the vlPAG (assessed using autoradiography) (Fullerton et al., 2021), and together, suggest an age-induced downregulation of PAG MOR in males matching a consistently lower level of MOR expression in females of any age. This apparent decrease in MOR expression may explain the reduced analgesic potency of morphine in females and aged males in our earlier work (Fullerton et al., 2021).

Here we report no impact of persistent inflammatory pain on vlPAG MOR binding potential, consistent with previous preclinical and clinical studies assessing the impact of chronic pain on MOR binding in the CNS. For example, patients with chronic fibromyalgia pain exhibit no changes in MOR binding in the PAG, despite having significant reductions in several brain regions, including the nucleus accumbens and the amygdala (Harris et al., 2007). Similarly, no change in PAG MOR binding was noted as a result of peripherally localized chronic pain generated from proximal nerve injury (Maarrawi et al., 2007).

Interestingly, centrally localized neuropathic pain (generated from supraspinal injury) was associated with a significant reduction of PAG MOR binding, suggesting that the origin of chronic pain within the CNS is critical in promoting alterations in neural MOR signaling (Maarrawi et al., 2007). Preclinical studies in rodents also suggest that changes in MOR induced by persistent pain are CNS site-specific. For example, persistent inflammatory pain induced by intraplantar CFA increased MOR binding in the dorsal root ganglion at 24 and 96 hours post-CFA, (Mousa et al., 2001; Zollner et al., 2003), with no change in MOR noted for the hypothalamus or spinal cord (Shaqura et al., 2004). Similarly, no change in PAG MOR availability or expression was observed following sciatic nerve injury, although reductions were noted for the insula, caudate putamen, and motor cortex (Thompson et al., 2018). Taken together, these results suggest that chronic pain alters MOR signaling in a CNS region-specific manner that is dependent on the site and modality of the pain.

### Impact of advanced age and sex on G-protein activation

Agonist binding at MOR activates Gαi/o-proteins and downstream signaling cascades which are critical for opioid-mediated hyperpolarization (Laugwitz et al., 1993; Connor and Christie, 1999; Koehl et al., 2018). In the present study, we found that adult males exhibit greater G-protein activation efficiency compared to aged males and adult and aged females, suggesting that the decreases in opioid potency observed in the aged males and the females are driven, in part, by attenuated G-protein-MOR coupling in the vlPAG. A significant impact of biological sex on G-protein potency (EC_50_) was also observed, with females and aged males showing a reduction in the potency of opioid-dependent G-protein activation, as evidenced by increased EC_50_ values. This reduction in vlPAG G-protein activation likely contributes to previous studies reporting that females are less sensitive to the analgesic effects of opioids (Kest et al., 2000; Loyd et al., 2008; Wang et al., 2014; Fullerton et al., 2018). Although the mechanism by which this change in potency occurs is not known, the nature of the GTPγS assay is suggestive. The assay uses membrane preparations without soluble signaling regulators like kinases present. Similarly, the GTPγS molecule itself is not hydrolyzable, and is thus not subject to regulation by GAPs like the RGS proteins. Considering these factors, the observed decrease in G protein activation potency may be due to the observed decrease in opioid receptor expression/availability in the binding studies above.

Interestingly, this was the only assay where an impact of persistent inflammatory pain was observed. Specifically, CFA treated rats had reduced EC_50_ values, regardless of age or sex. Reductions in Gα subunit expression have also been reported within both the rostral ventromedial medulla and spinal cord dorsal horn following intraplantar CFA in adult male rats (Wattiez et al., 2017). The mechanism whereby persistent inflammatory pain would impact G-protein activation, and not the other pharmacodynamics of MOR, is not known. Also unknown are the mechanisms by which advanced age and sex alter G-protein-MOR signaling. Previous studies have reported that advanced age results in a global downregulation of Gαi/o, most notably in the prefrontal cortex (Young et al., 1991; Alemany et al., 2007; de Oliveira et al., 2019), that is not associated with overall cell loss. Indeed, the Gαi/o subunit in particular appears to be highly susceptible to the aging process, with estimates of reduced expression as high as 65% in the frontal cortex, hippocampus, substantia nigra, and striatum (de Oliveira et al., 2019). Although age-induced changes in Gαi/o expression were not assessed specifically in PAG, a widespread reduction in Gαi/o as a function of advanced age would likely impact the PAG, thereby limiting MOR-G-protein coupling and reducing both G-protein activation efficiency and opioid potency in the aged rat.

Alternatively, an uncoupling between MOR and Gαi/o and/or a switch in Gα subunit from Gαi/o to Gαs cannot be ruled out (Gintzler and Chakrabarti, 2000, 2004, 2006). Indeed, a shift from Gαi/o to Gαs would similarly increase adenylate cyclase activity, resulting in reduced hyperpolarization and decreased morphine potency. Clearly, additional studies assessing the impact of advanced age on the expression of Gα isoforms are warranted (Lamberts et al., 2011).

The present studies used the high-affinity MOR agonist DAMGO for both the radioligand binding and GTPyS assay. As previous studies have reported that the pharmacodynamics of MOR may be agonist-dependent, this possibility cannot be ruled out and future studies should consider incorporating at least two agonists in all assays. Alternatively, a contribution by OPRM1 splice variants induced as a function of age may also be present (Pan et al., 2005; Alvarez et al., 2002; Narayan et al., 2021). Indeed, specific alternative MOR isoforms are known to promote ligand-dependent regulation of opioid signaling (Pasternak et al., 2004; Oldfield et al., 2008). MOR-1B, a family of MOR isoforms expressed in the brain, are indistinguishable from MOR-1 at the extracellular surface and the agonist binding pocket but elicit alternative signaling properties that impact opioid-induced cell hyperpolarization, and dysregulation of these MOR splice variants has been implicated in advanced age (Latorre and Harries, 2017; Li et al., 2017; Wang et al., 2018). There are also reports of sex differences in the expression of MOR splice variants in the PAG, with male rodents expressing higher levels of MOR-1B5, and females expressing higher levels of MOR-1B1 and MOR-1B2 (Liu et al., 2018). The unique conformation of MOR isoforms and subsequent alterations in receptor phosphorylation would directly influence the G-proteins to which MORs couple, potentially resulting in biased agonism (Verzillo et al., 2014; Abrimian et al., 2021).

### Impact of advanced age and sex on MOR Phosphorylation

The results of the radioligand binding and GTPγS assays suggested that aged and female rats exhibit decreased receptor expression and activation potency. It has been previously shown that MOR can be desensitized by phosphorylation in its basal state, thereby limiting agonist activation (Zhang et al., 1996; Yu et al., 1997; Groer et al., 2011). Thus, we tested the hypothesis that the observed reductions in G-protein activation efficiency were due to age- and sex-induced changes in basal MOR phosphorylation. Although aged rats expressed higher levels of MOR phosphorylation at serine-375 compared to the adults, this result was not significant, suggesting that the reductions in activation potency observed in the vlPAG of aged and female rats is likely driven by an alternative mechanism.

MOR desensitization is mediated by G protein-coupled receptor kinase (GRK)-dependent phosphorylation of the receptor, a mechanism by which β-arrestins bind to MOR, resulting in endocytosis (J. Zhang et al., 1998; Schulz et al., 2004; Dang et al., 2009). Although our results suggest no impact of advanced age or sex on GRK-mediated MOR phosphorylation, MOR signaling is also desensitized via phosphorylation through the extracellular signal-regulated kinase 1 and 2 (ERK1/2) pathway (Dang et al., 2009). ERK1/2 phosphorylation stimulates the activity of Gα-interacting protein (GAIP), an RGS protein that acts as a GTPase activator to reduce opioid signaling at the level of G-protein activation (Ogier-Denis et al., 2000). Indeed, pharmacological inhibition of ERK1/2 phosphorylation in a rat model leads to improved morphine analgesia (Popiolek-Barczyk et al., 2014; Melkes et al., 2020). Thus, the age- and sex-induced changes in morphine potency may be driven in part by age-induced hyper-phosphorylation of ERK1/2 and result in the downregulation of G-protein signaling.

### Impact of advanced age and sex on cAMP inhibition

Agonist binding at MOR elicits a conformational change in the receptor that allows the G-protein α and βγ subunits to interact with downstream effectors. Notably, the Gα subunit binds to adenylate cyclase (AC) and inhibits the conversion of adenosine triphosphate (ATP) to cAMP, thus limiting the activation of cAMP-dependent protein kinase (PKA) and ultimately inducing higher levels of hyperpolarization (Christie, 2008; Santhappan et al., 2015). In the present study, aged males and adult and aged females exhibited significantly lower levels of DAMGO-induced cAMP inhibition compared to adult males, suggesting an attenuated activity of the α subunit at the level of AC, or a weakened relationship between AC and cAMP. This effect could easily be due to the observed decreases in MOR expression and activation potency observed above. This attenuated cAMP inhibition in the aged and female rats likely contributes to the observed reduction in opioid potency and may also be due, in part, to decreased basal cAMP levels in the aged brain. Reductions in cAMP have been reported in several rodent brain regions including the cortex, thalamus, hypothalamus, and midbrain (Puri and Volicer, 1981; Titus et al., 2013; Kelly, 2018). Additional evidence suggests that aged rodents elicit widespread reductions in ATP (Błaszczyk, 2020), which is required for cAMP synthesis. Not surprisingly, previous studies assessing the impact of advanced age on cAMP expression and signaling were conducted exclusively in male rodents, so it is not known whether advanced age impacts neural cAMP levels in a sex-specific manner.

Morphine binding to MOR results in the activation of Gai/o that primarily inhibits the activity of AC, thereby reducing the ability to generate cAMP; in parallel, the βγ subunits activate G-protein inwardly rectifying potassium (GIRK) channels to induce hyperpolarization and decrease excitability (Law et al., 2000; Nobles et al., 2005; Yudin and Rohacs, 2018). Advanced age is concomitant with increased levels of neuroinflammation and greater production of reactive oxygen species (ROS), both of which have been implicated in age-induced potassium channel dysfunction (Sparkman and Johnson, 2008; Sesti, 2016; Lieberman et al., 2020). Consistent with this, aged Sprague Dawley rats (26-30 mos) exhibit reduced potassium currents in hippocampal neurons (Alshuaib et al., 2001). Further experimentation assessing the impact of advanced age and sex on GIRK channel-mediated hyperpolarization, in addition to cAMP-mediated hyperpolarization, will provide a more detailed description of the impact of advanced age and sex on downstream opioid signaling in the PAG.

### Impact of advanced age and sex on RGS protein levels

G-protein activation by MOR results in the exchange of GDP for GTP (Senese et al., 2020). GTP-bound Gα proteins are inactivated by hydrolysis of the GTP back to GDP, catalyzed by the intrinsic GTPase activity of Gα proteins. This inactivation mechanism is modulated by the activity of Regulator of G-protein signaling (RGS) proteins. RGS proteins reduce the magnitude of GPCR signaling via enhancement of GTPase activity of the α subunit of the G-protein complex. By promoting the hydrolysis of the alpha-bound GTP during the active state, RGS proteins hasten the return of the α subunit to the GDP-bound inactive state (Roman and Traynor, 2011). RGS proteins play a critical role in negatively modulating opioid signaling, as morphine analgesia is increased in male mice lacking RGS9-2 (Garzón et al., 2001; Zachariou et al., 2003) and overexpression of RGS4 attenuates MOR signaling in reconstituted MORs in vitro (Ippolito et al., 2002). In the present study, we observed increased expression of RGS4 and RGS9-2 in the vlPAG of aged rats compared to adults, suggesting greater GTPase activity and thus reduced G-protein signaling in the aged. Similar results were observed when the analysis was limited to MOR+ cells, indicating that opioid-induced G-protein signaling is subjected to greater negative regulation in the vlPAG of the aged rat. These results are consistent with Kim et al. (2005) who reported higher RGS9-2 protein levels in the PAG of 1-year old male rats compared to 3-week-old rats (Kim et al., 2005). In humans, advanced age is associated with increased RGS4 expression in the prefrontal cortex (Rivero et al., 2010), suggesting that the age-induced increase in RGS4 in our experiment may not be specific to the vlPAG. These results provide a mechanism by which advanced age could decrease analgesic potency and downstream cAMP signaling, although they do not provide a mechanism by which adult females demonstrate lower potency and signaling than adult males.

## Summary and Conclusions

The present studies are the first to show that sex and advanced age leads to an attenuation in vlPAG opioid signaling compared to adult male rats. Taken together with our previous findings, these results suggest that age- and sex-induced reductions in vlPAG MOR expression and binding, combined with attenuated downstream MOR signaling, contribute to the diminished opioid potency reported in aged and female rats (Fullerton et al., 2021). The results of our analyses demonstrate that aged and female rats exhibit reductions in MOR expression/availability, G-protein activation, and cAMP inhibition, and increased G-protein regulation by RGS proteins, each of which provides potential therapeutic targets for improved pain management in the elderly. The nature of the results observed also suggest potential mechanisms of action for sex and age to impact analgesic potency. First, the normal ligand affinity (Kd) and lack of receptor phosphorylation suggests that unit receptor performance is normal. When combined with the observed decrease in expression/availability (Bmax), this suggests that one major mechanism is a decrease in receptor expression instead of a change in unit receptor performance. When considering the G protein coupling analysis, since soluble regulators (like RGS proteins) are absent in this assay and the ^35^S-GTPγS is non-hydrolyzable, the decrease in G protein coupling potency is likely due to decreased receptor availability. Downstream of the G proteins, we also observed a decrease in cAMP inhibition. This downstream effect could be due to the observed increase in RGS expression, decreasing G protein activity, leading to less cAMP inhibition. However, RGS expression was not increased in adult females. Overall then our results suggest two major complementary mechanisms; 1) decreased receptor expression/availability in adult females and aged males and females, and 2) increased RGS expression in aged males and females. Further investigation could also uncover additional complementary sex- and age-related mechanisms that help explain why opioids show lower analgesic potency in females and the aged.

## Author Contributions

AZM and EFF designed the research and wrote the paper. MK, JMS, and LJY aided in the design of the research. EFF, MK, and JMS performed the research, EFF, AZM, and JMS analyzed the data.

